# Living electronic transistors with tunable conductivity

**DOI:** 10.64898/2026.09.01.748419

**Authors:** Joshua T. Atkinson, Marko S. Chavez, Christina M. Niman, Fengjie Zhao, Mi So Hwang, Nir Sukenik, Jeffrey A. Gralnick, James Q. Boedicker, Mohamed Y. El-Naggar

## Abstract

Electroactive bacteria, like *Shewanella oneidensis*, can couple the oxidation of organic electron donors to the reduction of external conductive surfaces, such as minerals and electrodes, by utilizing multiheme cytochromes to carry charge from within the cell to external surfaces. Additionally, multiheme cytochromes facilitate gateable, long-distance (micrometer-scale) redox conduction along the outer membrane and across multiple cells bridging electrodes. While electroactive microbes are being used to develop bioelectrochemical devices, there have been limited efforts to use synthetic biology to exert additional control over microbes serving as device components. Thus, this work implements an optogenetic biofilm patterning gene circuit and a small molecule sensor in *S. oneidensis* to simultaneously control cell deposition and cytochrome expression. This allows for photolithographic patterning of biofilms possessing tunable electrical properties controlled with small molecules. This system demonstrates tunable electrochemical activity, redox conduction, intrinsic biofilm conductivity, and negative differential transconductance as a function of cytochrome expression. Additionally, temperature-dependent measurements of this tunable biofilm conduction reveal changes in activation energy as a function of cytochrome expression. Through this combination of synthetic biology and electrochemistry, simultaneous control over biofilm geometry and conductivity sheds light on fundamental microbial electron transport processes, and it enables the construction of living electronic devices.

## INTRODUCTION

Electroactive microorganisms can couple the oxidation of reduced substrates to the reduction of external solid electron acceptors, such as minerals and conductive electrode surfaces, in a process called extracellular electron transport (EET) ^1–3^. To perform EET, many electroactive organisms, including *Shewanella oneidensis,* utilize a series of multiheme cytochromes capable of carrying metabolic electrons across the cell envelope to external surfaces, which can be utilized for biotic-abiotic electrical interfacing with electrochemical devices^3–8^. Additionally, these multiheme cytochromes facilitate micrometer-scale, thermally activated, redox conduction along the outer membrane and across multiple cells within biofilms bridging electrodes via cytochrome-cytochrome and cell-cell electron transport^9–12^.

Functioning as living catalysts, electroactive biofilms have found applications as components of biohybrid devices for renewable energy generation in microbial fuel cells, for chemical production in microbial electrosynthesis cells, and for wastewater treatment in microbial electrolysis cells^13^. Beyond these applications, electroactive biofilms present an ideal platform for developing living electronic technologies. Living electronics are novel electronic devices which seamlessly interface living and non-living systems while possessing the flexibility, self-assembly, self-repairing, and sensing capabilities of biological systems as well as the tunability, conductivity, and electrical readout capabilities of traditional solid-state devices^14–18^. To realize this new generation of technology, we need methods to control microbial-based systems similar to those used in solid-state electronics fabrication, such as photolithographic patterning and conductivity tuning via ion doping. To implement such controls in electroactive bacteria, we turn to the genetic engineering toolbox of synthetic biology.

Synthetic biology techniques have enabled the control of bacterial behavior by modulating transcription, translation, and protein activity in response to diverse environmental stimuli including light^19^, temperature^20^, chemicals^21^, and electric potentials^22^. Additionally, these techniques have been used to confer electroactivity to the non-electroactive model bacterium *Escherichia coli*, and to control the current production of native electroactive bacteria such as *S. oneidensis* and *Geobacter sulfurreducens* ^15,23–33^. In one recent work, light was used to spatially pattern electroactive biofilms, and in separate studies, it was used to control EET in electroactive biofilms^12,34,35^. These works open the door for photolithographic approaches to be used in controlling living electronic device fabrication. However, to date, no study has demonstrated combined control over both the spatial geometry and the electrical properties of bacterial biofilms.

In this work, we implemented an optogenetic biofilm patterning gene circuit in the electroactive organism *S. oneidensis* alongside a chemogenetic gene circuit to control expression of cell surface cytochromes. This allowed us to pattern biofilms onto electrode surfaces with light and then tune their electrical properties using a chemical inducer to create living chemiresistors that change conductivity in response to added chemicals. We first examined the EET capabilities of patterned *S. oneidensis* on planar electrodes as a function of cytochrome content. Then, using interdigitated electrodes (IDEs), we performed electrochemical gating measurements on patterned biofilms to determine their conductivity while controlling cytochrome concentration to demonstrate tunability in the electrical properties of these engineered living materials. Additionally, temperature-dependent conduction measurements were performed at each cytochrome expression level to determine how the activation energy associated with the redox conduction mechanism changes as a function of cytochrome content. From these results, we observed a modest decreasing trend in activation energy as cytochrome expression was increased, suggesting a change in the mechanism of electron transport as a function of cytochrome content on the cell surface. Finally, we show that these living electronic films behave like transistors possessing negative differential transconductance (NDT), typically used in multivalued logic devices, by extracting NDT device figures of merit and differential transconductance from electrochemical gating data. Taken together, our results demonstrate how synthetic biology enables control over microbial-based biomaterials similar to what is used to manipulate solid-state materials in semiconductor device fabrication.

## METHODS

### Plasmids and Strains

A list of all plasmids and strains used in this study can be found in Table S1 and Table S2, respectively. The patterning plasmid was pDawn-CdrAB^21^

(**Figure 1A**). pMtrCAB was generated by PCR amplifying *mtrCAB* from *S. oneidensis* genomic DNA and were assembled into a vanillic acid inducible vector pAJM.773^25^ using Golden Gate cloning^36^. RBS sequences were designed using the RBS calculator v2.1.1 (De Novo DNA)^37^ and were synthesized as oligos containing overhangs that were phosphorylated, annealed, and inserted between the P_van_ promoter and *mtrC* using Golden Gate cloning.

**Figure 1.**
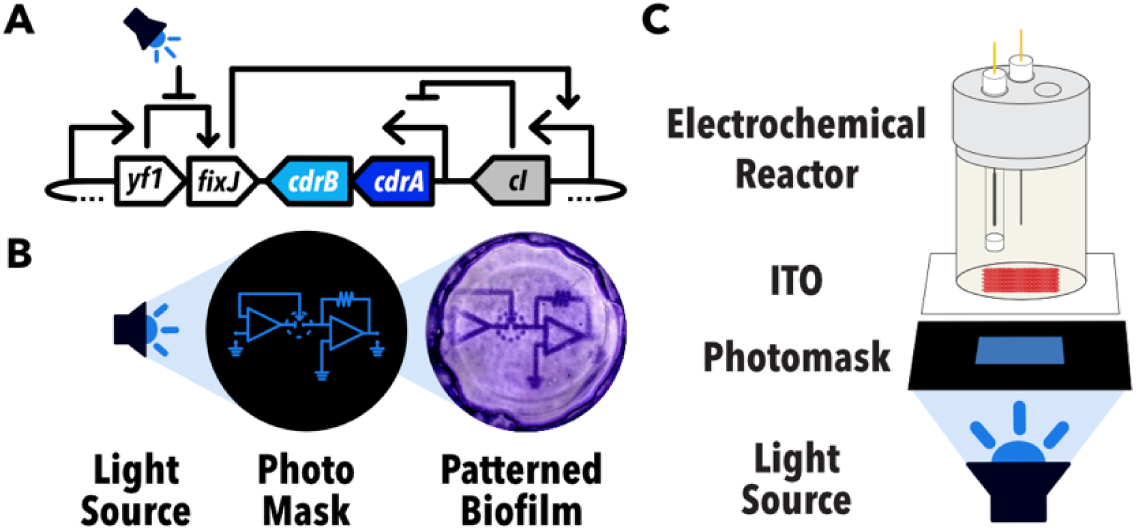
Optogenetic patterning of biofilms. **(A)** Gene circuit for controlling expression of biofilm formation proteins (CdrAB). Blue light inhibits phosphorylation of FixJ by Yf1 leading a drop in expression of the repressor cI resulting in expression of CdrAB. **(B)** Biofilms can be patterned onto transparent surfaces using a light-source and a photo mask as shown by the crystal violet-stained biofilm after patterning. **(C)** Photolithography into electrochemical reactors using transparent Indium Tin oxide (ITO) electrodes.

### Growth conditions

*S. oneidensis* strains were cultivated in lysogeny broth (LB) or minimal medium (MM)^38^ (15.1 g/L PIPES buffer, 3.4 g/L NaOH, 1.5 g/L NH_4_Cl, 0.1 g/L KCl, 0.6 g/L NaH_2_PO_4_·H_2_O, 3.36 g/L 60% (w/w) sodium DL-lactate, 1 mL mineral solution, 1 mL vitamin solution^39^, 10 mL amino acid solution, pH 7.0) at 30 °C, 250 rpm (25 mm throw). The mineral solution consists of 15 g/L nitrilotriacetic acid (dissolved with NaOH to pH 8.0), 30 g/L MgSO_4_·7H_2_O, 5 g/L MnSO_4_·H_2_O, 10 g/L NaCl, 1 g/L FeSO_4_·7H_2_O, 1 g/L CaCl_2_·2H_2_O, 1 g/L CoCl_2_·6H_2_O, 1.3 g/L ZnCl_2_, 0.1 g/L CuSO_4_·5H_2_O, 0.1 g/L AlK(SO_4_)_2_·12H_2_O, 0.1 g/L H_3_BO_3_, 0.25 g/L Na_2_MoO_4_·2H_2_O, 0.24 g/L NiCl_2_·6H_2_O and 0.25 g/L Na_2_WO_4_·2H_2_O. The vitamin solution consists of 0.02 g/L D-biotin, 0.02 g/L folic acid, 0.1 g/L pyridoxine HCl, 0.05 g/L riboflavin, 0.05 g/L thiamine HCl·H_2_O, 0.05 g/L nicotinic acid, 0.05 g/L calcium D-pantothenate, 0.001 g/L cyanocobalamin (B12), 0.05 g/L p-aminobenzoic acid, and 0.05 g/L thioctic acid. The amino acid solution consists of 2 g/L L-glutamic acid, 2 g/L L-arginine, and 2 g/L DL-serine. When necessary, media were supplemented with kanamycin (Kan, 50 μg/mL) and streptomycin (Strep, 100 μg/mL) for plasmid selection.

### Relative assessment of cytochrome expression

Relative cytochrome expression was assessed using either heme staining (Figure 2) or enhanced chemiluminesence (Figure S1). For heme staining detection, cultures of *S. oneidensis* ΔMTR, which lacks genes encoding outer-membrane and periplasmic cytochromes essential for EET, containing either pMtrCAB-low or pEmpty were grown in MM supplemented with 50 μg/mL kanamycin and varying amounts of vanillic acid at 30°C with shaking at 250 rpm (25 mm throw) for 20 h. Cultures were then adjusted to an optical density of 0.4 in fresh MM, mixed 1:1 with Laemmli buffer, incubated for 10 min at 100°C, and ran on an SDS-PAGE gel. In-gel peroxidase activity of cytochromes was detected by staining using the chromogenic substrate 3,3’,5,5’-tetramethylbenzidine (TMBZ). Gels were soaked with 50 mL of TMBZ stain solution (2 parts of 6.3 mM TMBZ dissolved in 100% methanol and 1 part of 250 mM sodium acetate, pH = 5.0) for 1 h, followed by development with 1.5 mL of 30 % hydrogen peroxide for 30 min. Gels were then imaged using a flatbed scanner. For enhanced chemiluminescence detection^39^, cultures of *S. oneidensis* ΔMTR containing pMtrCAB-low, pMtrCAB-med, pMtrCAB-high, or pMtrCAB-highest were grown overnight in LB supplemented with 50 μg/mL kanamycin at 30 °C with shaking at 250 rpm (25 mm throw). Overnight cultures were normalized to an OD600 of 0.05 and subcultured in 5 mL LB supplemented with either 0 μM or 25 μM vanillic acid and grown at 30 °C with shaking at 250 rpm (25 mm throw). Cells were normalized to OD_600_ of 10 mixed 1:1 with Laemmli buffer, incubated for 10 min at 100°C, and ran on an SDS-PAGE gel. The gel was transferred to a nitrocellulose membrane using the Trans-Blot Turbo Transfer System (Bio-Rad) and enhanced chemiluminescence detection was performed using Clarity^TM^ Western ECL Substrate (Bio-Rad).

**Figure 2.**
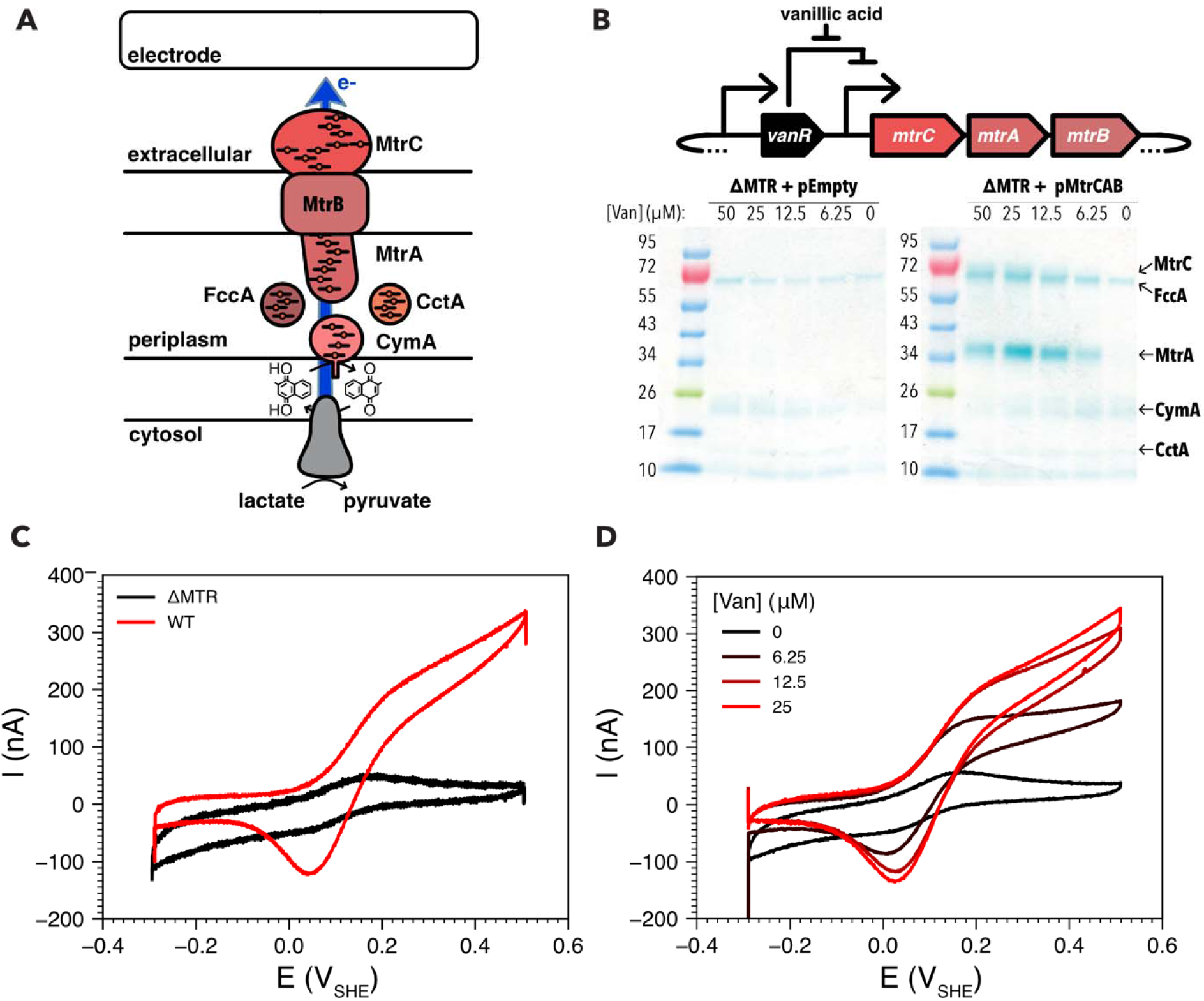
Tunable cytochrome production. **(A)** Extracellular electron transport chain from *S. oneidensis* that forms a conduit to move electrons from lactate dehydrogenase in the cytosol, to the quinol oxidase CymA in the inner membrane, to the multiheme cytochrome MtrA in the periplasm and ultimately to multiheme cytochrome MtrC on the cell surface through the MtrB porin. **(B)** (Top) Gene circuit used for controlling expression of the cytochrome-porin complex using vanillic acid. (Bottom) A heme-stained SDS-PAGE gel of cultures of a multiheme cytochrome mutant (ΔMTR) with the gene circuit (pMtrCAB-low) or with an empty vector (pEmpty) with varying concentrations of vanillic acid. MtrA and MtrC bands increase with vanillic acid while FccA, a genomically-encoded cytochrome is present under all conditions as a loading control. **(C)** Turnover cyclic voltammetry of WT and multiheme cytochrome mutant (ΔMTR) in the presence of the electron donor lactate. WT generates a catalytic current at potentials >0.1 V_SHE_, while ΔMTR does not. **(D)** Turnover cyclic voltammetry of ΔMTR containing the genetic circuit in the presence of various concentrations of vanillic acid. For each condition, the data shown are representative traces of a single replicate from a triplicate set.

### SpyCatcher-GFP expression

*E. coli* BL21(DE3) cells transformed with pSpyCatcher003-sfGFP (Addgene #133449) were grown overnight in 5 mL of LB supplemented with 100 μg/mL carbenicillin at 37°C with shaking at 250 rpm (25 mm throw). Overnight culture was subcultured at a 1:100 inoculum in 600 mL of LB. Once the cells reached an optical density at 600 nm (OD_600_) of 0.6, isopropyl β-D-1- thiogalactopyranoside was added to a final concentration of 0.5 mM to induce protein expression. After 18 h of induction at 15°C with shaking at 150 rpm, cells were harvested by centrifugation at 3000 rcf for 20 min at 4°C. The pellet was frozen at −80°C for later protein purification.

### Purification of SpyCatcher-GFP

His-tagged SpyCatcher-GFP (SC-GFP) was purified using Ni-NTA affinity chromatography. The cell pellet (1.5 g wet) was resuspended in 12 mL of equilibration buffer containing 43 mM Na_2_HPO_4,_ 7 mM NaH_2_PO_4_, 300 mM NaCl and supplemented with 100 μL of 10x Halt™ Protease Inhibitor Cocktail, EDTA-Free (Thermo Scientific), 2 μL of benzonase nuclease (Millipore Sigma), and 2 mg chicken egg white lysozyme (GoldBio). Cells were lysed by sonication (QSonica Q500D, cup horn, amplitude 20%, pulse: 10s ON/20s OFF, 10 cycles, time: 4min), and the lysate was clarified by centrifugation at 7000 rcf for 15 min at 4°C. 12 mL of lysate was applied to gravity flow columns (Marvelgent Biosciences) containing 2 mL of PROTEINDEX^TM^ HiBond^TM^ Ni-NTA Agarose resin (Marvelgent Biosciences). The lysate was washed with 12 mL of equilibration buffer containing 20 mM imidazole (GoldBio) and eluted using 5 mL of equilibration buffer containing 250 mM imidazole. The fractions were collected and analyzed by SDS-PAGE to determine relative purity. Protein samples were dialyzed into PBS with Slide-A-Lyzer™ G3 Dialysis Cassettes, 10K MWCO (Thermo Scientific), aliquoted and stored at −80 °C till later use. Protein concentration was determined using Pierce™ Bradford Plus Protein Assay Kit (Thermo Scientific) and absorbance was measured at 595 nm.

### Cell-surface cytochrome quantification using SpyCatcher-GFP labelling

Labelling experiments were conducted using *S. oneidensis* ΔMTR containing pMtrCAB-low-ST, which encodes MtrC containing a SpyTag peptide insertion following amino acid 245. Cell cultures were grown overnight in 5 mL LB supplemented with 50 μg/mL kanamycin at 30 °C with shaking at 250 rpm (25 mm throw). The culture was normalizing to OD_600_ of 0.05 and subcultured in MM supplemented with with 50 μg/mL kanamycin and varying concentrations of vanillic acid (0 μM, 1.5625 μM, 3.125 μM, 6.25 μM, 12.5 μM, and 25 μM). The subculture was grown overnight at 30 °C while being shaken at 250 rpm (25 mm throw). SC-GFP was determined to be saturating at 20 μM and was used for all subsequent labelling experiments. For cell surface MtrC labelling, overnight cultured cells were normalized to OD_600_ of 1.0 in 500 μL of PBS in a 1.5 mL tube and 20 µM of purified SC-GFP was added. The mix was allowed to incubate for 1 h at room temperature with 150 rpm shaking. Cells were washed with PBS two times to remove unbound SC-GFP before resuspension in 500 μL PBS. A 200 μL aliquot of the suspension was transferred to a black 96 well plate (Corning, 3603) and GFP fluorescence was measured with a plate reader (Agilent BioTek Synergy H1; excitation: 485 nm, emission: 510nm). A SC-GFP standard curve was generated by measuring the fluorescence of *S. oneidensis* ΔMTR without pMtrCAB-low-ST incubated with 0-2 μM SC-GFP. The SC-GFP/cell was then calculated using the standard curve and the colony-forming units for the cell culture.

### Transparent-bottom bioreactor construction

To observe the surface coverage and thickness of patterned biofilms on bioreactor working electrodes (WEs) *in situ*, transparent-bottom bioreactors were constructed. Planar, 22 mm by 40 mm indium tin oxide (ITO) coated glass coverslips (planar-ITO, Prod no. 06494-AB, Structure Probe, Inc.) or custom ITO IDEs (designed in house and fabricated via foundry service) were used as the bioreactor base. Before use, the ITO coated coverslips and the custom IDEs were rinsed with acetone, isopropanol, and then with DI water, and then dried with N_2_. Thin copper wires (Prod no. 1227, TCS 20’ 32 gauge) were electrically connected to the WEs with silver paint (Prod no. 16035, TED PELLA, Inc.). To mechanically strengthen the wire-electrode connections, they were covered with epoxy (Gorilla Glue Co.). To function as the body of the bioreactors, glass tubes (2.5 cm tall with a 19 mm and 22 mm inner and outer diameter, respectively) were adhered overtop of the WEs with siliconized sealant (DAP Kwik Seal Ultra Premium Siliconized Sealant). Custom PEEK plastic lids were used to hold Pt wire counter electrodes (CEs) and 1 M KCl Ag/AgCl reference electrodes (REs) (CHI111P, CH Instruments, Inc.). All potentials reported in this manuscript are corrected to Standard Hydrogen Electrode by adding +0.235 V to correct for the 1 M KCl electrolyte in the Ag/AgCl reference.

### Microfabrication of custom interdigitated array electrodes

Custom indium tin oxide (ITO) interdigitated electrodes (IDEs) were fabricated by coating glass with ITO to form 15 μm insulating gaps between adjacent, interdigitated electrode fingers. These devices were designed in house and were fabricated by the University of California, San Diego Nano3 cleanroom foundry service using standard photolithography techniques. The fingers for each device were 12 mm long and 10 μm wide. There were 200 electrode finger pairs for the 15 μm gap IDEs, with a total WE area of 0.48 cm^2^ (**Figure 3B**, Zenodo Dataset - 10.5281/zenodo.21959616). The fabrication process is briefly outlined as follows. 100 mm diameter and 500 mm thick BK-7 glass wafers (University Wafer, Inc.) were used as the electrode substrate. Solvent cleaned wafers were coated with photoresist (PR) and then a laser writer was used to project multiple IDE patterns, side by side, onto the PR with UV light. PR developer was then used to remove the light-exposed PR and then 300 nm of ITO was sputter coated onto the wafers. Solvents were used to remove the excess ITO and PR, leaving just the electrode pattern. To improve the conductivity of the electrodes, the wafers were baked in an N_2_ furnace at 400 °C for 1 h^40^. As a final step, the wafers were coated in PR and then diced into multiple 24 mm by 60 mm chips, with each chip containing one ITO IDE pattern. It must also be noted that since the interdigitated area of these devices (including fingers and gaps) was 1.2 x 1 cm^2^, the effective biofilm pattern sizes used in our electrochemical gating measurement are 1 mm smaller lengthwise than the projected patterns.

**Figure 3.**
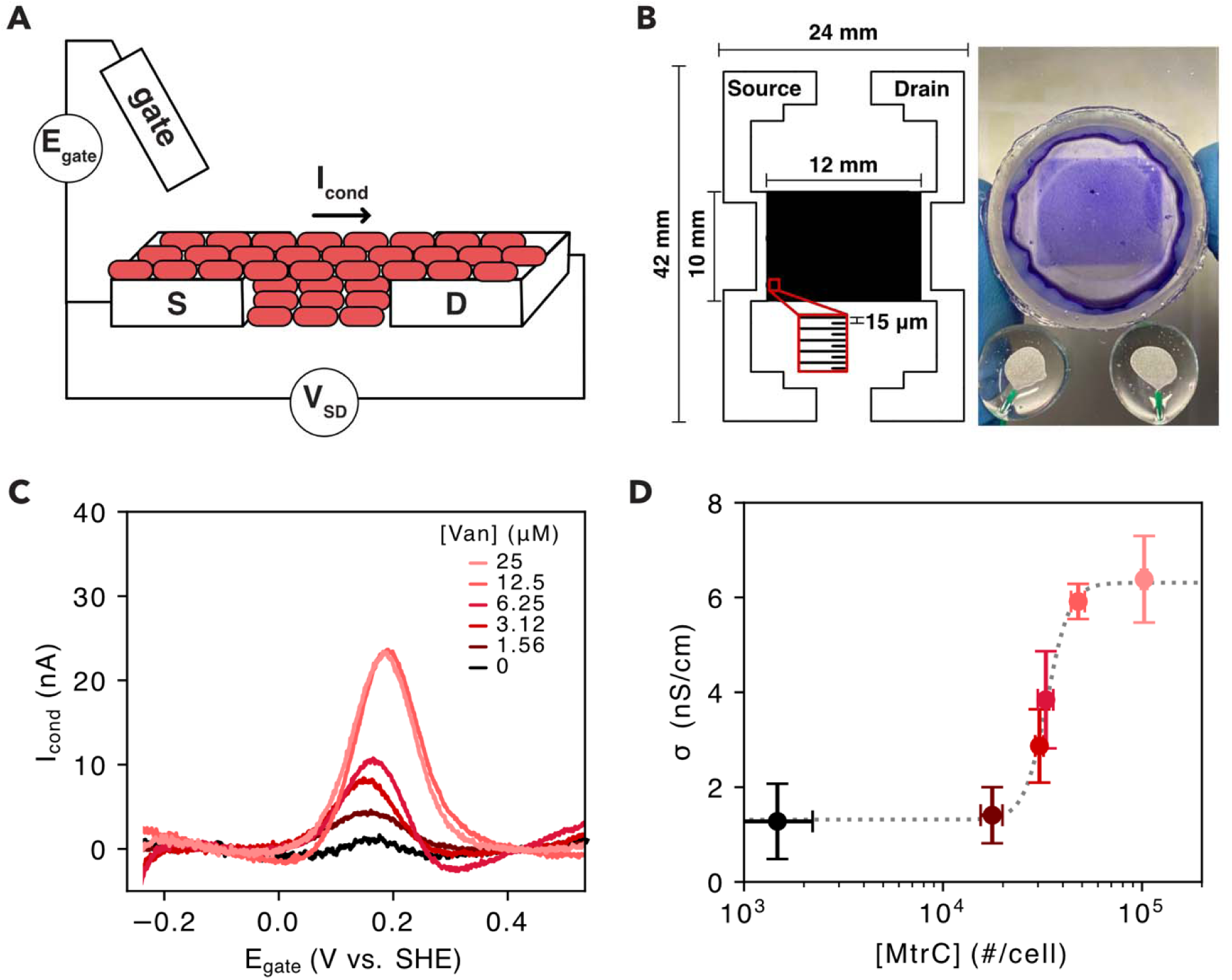
Electrochemical gating of electron transport through bacterial biofilms with varying cytochrome content. **(A)** Schematic of electrochemical gating devices with biofilms (red) spanning source (S) and drain (D) electrodes that are controlled using a gate electrode to apply a gate potential (E_gate_) and a source-drain voltage (V_SD_) to drive a conduction current (I_cond_). **(B)** Left - Schematic of interdigitated ITO electrode arrays used for measuring biofilm conductivity. Right - Photograph of crystal violet stained biofilm following biofilm photolithography**. (C)** Conduction current (I_cond_) vs gate potential (E_gate_) from electrochemical gating measurements of patterned biofilms of the ΔMTR containing the pMtrCAB-low genetic circuit and in the presence of various concentrations of vanillic acid. **(D)** Biofilm conductivity relative to the cytochrome abundance per cell when cells are induced with varying concentrations of vanillic acid. Circles and error bars represent the mean and standard deviation, respectively, of biological replicates (n=3-5 per condition). The dotted line represents a Hill function fit to the data to calculate the half-maximal cytochrome concentrations.

### Light-patterned biofilm formation workflow

Cells were inoculated, from frozen glycerol stock, into 5 mL of sterile LB and incubated for 21 h, in the dark, at 30^°^C with shaking at 250 rpm (25 mm throw). The LB was supplemented with 100 μg/mL of streptomycin and 50 μg/mL of kanamycin to select for the light-patterning plasmid and the cytochrome content tuning plasmid, respectively. MM, supplemented with 40 mM sodium fumarate and the necessary antibiotics, was inoculated with the LB culture to a final OD_600_ of 0.01. Varying concentrations of vanillic acid were also added at this step depending on the desired cytochrome expression. Then, 2 mL of MM was added to the transparent bottom bioreactors, which were sealed with gas permeable flask seals (Prod no. 899421, Thomson Ultra Yield 0.2 m). For light-patterning (**Figure 1B-C**), the bioreactors were incubated for 21 h on top of a photomask, light dispersal sheet, and an LED matrix array (_peak_ = 468 nm, Prod no. KWM-R30881XBB, Luckylight) set to an irradiance of 110 μW cm^-2^ using a microcontroller (Adafruit Feather RP2040) as measured using an optical power meter (S120VC photodiode sensor and PM100USB interface, Thorlabs). During light-patterning, the bioreactors acted solely as culturing vessels and not as electrochemical cells.

Following patterning, the media in the bioreactors was exchanged for 1 mL of sterile PBS. The bioreactors were then lightly shaken on an orbital shaker for 60 s. This step was repeated twice. The bioreactors were then brought into the anaerobic chamber (Bactron 300, Sheldon Manufacturing, Inc.) with an N_2_/H_2_ (95:5) atmosphere where the PBS was exchanged for anaerobic MM. For turnover cyclic voltammetry experiments on planar-ITO electrodes, turnover MM without fumarate, or vitamins was used (**Figure 2C- D**). For electrochemical gating experiments on ITO-IDEs, non-turnover MM without lactate, fumarate, or vitamins was used (**Figure 3C-D**).

### Measurements of electrochemical activity and biofilm conduction

All electrochemical measurements were performed in an anaerobic chamber with an N_2_/H_2_ (95:5) atmosphere. For all experiments, sterile medium electrochemical measurements were performed before reactors were used in biofilm patterning. For the planar-ITO reactors, cyclic voltammetry (CV) measurements were performed by scanning the WE potential from -265 mV_SHE_ to 535 mV_SHE_ at 1 mV sec^-1^ using a four channel potentiostat (Squidstat Prime, Admiral Instruments). For ITO-IDE reactors, electrochemical gating measurements were performed using a 6 channel potentiostat (VSP300, Bio-Logic) to sweep the WE potentials at 1 mV sec^-1^ from -265 mV_SHE_ to 535 mV_SHE_ with a fixed gating offset *V*_SD_ = 20 mV held between the two WEs. Here, *V*_SD_ = *E*_D_ – *E*_S_, where *E*_D_ and *E*_S_ are the potentials at each of the IDE WEs. Data are reported relative to *E*_gate_ which is the average of *E*_D_ and *E*_S_. To determine *I*_cond_, we assumed that equal and opposite sign *I*_cond_ plus background currents of the same sign were measured at each IDE WE^12^. Thus, *I*_cond_ could be calculated by subtracting the drain and source currents and dividing by two, (*I*_D_ – *I*_S_)/2. Three cycles were performed for both the CV and gating scans and only data from the third cycle is presented in this manuscript. Electrochemical gating measurements at each vanillic acid concentration were performed at least in triplicate, with representative *I*_cond_ vs *E*_gate_ plots shown in **Figure 3**. Biofilm conductivity was derived using Ohm’s law and a conformal mapping technique using Equations 1-3 as previously described^12^.The term a is the half of the width of a finger gap and the term g is the biofilm thickness.

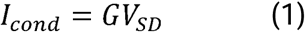

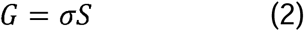

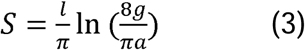

### *In situ* confocal microscopy, biofilm thickness analysis, and crystal violet staining of patterned biofilms

After electrochemical measurements, the reactor medium was discarded, and 2 mL of MM containing 0.25 μg mL^-1^ FM 4-64FX membrane dye (Invitrogen Cat. No. F34653) was added to the reactors. The biofilms were then incubated in the dark for 30 min to allow membrane staining. The medium was subsequently replaced with 2 mL of 3% glutaraldehyde solution, and the biofilms were fixed at 4^°^C overnight. The samples were then washed three times with 2 mL of PBS. A Zeiss LSM 880 inverted microscope equipped with an argon laser and either a 20× dry objective or a 40× water-immersion objective was used to image the biofilms. Imaris software was used to process the confocal image stacks and generate three-dimensional reconstructions and cross-sectional images of the biofilms. Biofilm thickness was determined by analyzing the cross-sectional confocal images using MATLAB, as previously described ^12,24^. Crystal violet staining was performed after confocal imaging by adding 1 mL of 0.1% crystal violet solution for 10 min, followed by three rinses with 2 mL of PBS to visualize the patterned biofilms^12,35^.

## RESULTS AND DISCUSSION

### Synthetic biology enables programable deposition and electroactivity of biofilms

To control biomass deposition and spatially define the area in which multicellular biofilms formed on electrode surfaces, we leveraged an optogenetic approach that was recently shown to enable photolithographic printing of *S. oneidensis* biofilms^26^. *S. oneidensis* was transformed with the plasmid pDawn-CdrAB, encoding cell aggregation proteins CdrAB from *Pseudomonas aeruginosa* under the control of the blue-light inducible transcriptional promoter (P_Dawn_) (**Figure 1A**). This synthetic gene circuit enables photolithographic patterning of biofilms (**Figure 1B**) directly onto electrodes within electrochemical cells when using a light source and photomasks (**Figure 1C**). Throughout this study, we utilized this plasmid system to precisely deposit biomass on electrodes regardless of the cytochrome content of the strain being deposited.

To complement this spatial control of biofilm formation, we developed a chemogenetic system for controlling the concentration of the MtrCAB cytochrome-porin complex on the cell surface (**Figure 2A)**. Initially, a series of plasmids (pMtrCAB-low, - medium, -high, -highest) were built to control the expression of MtrCAB in *S. oneidensis* using a vanillic acid-inducible promoter (P_vanR_)^37^. Each plasmid was designed to have an initial ribosomal binding sites (RBS) for *mtrC* in the *mtrCAB* operon with varying translation initiation efficiencies (*e.g.,* 800, 1100, 3500, 7500 au)^40^. To determine the RBS best suited for controlling cytochrome expression, strains containing each plasmid were screened for cytochrome production with and without vanillic acid (**Figure S1**). After comparing constructs, it was determined that pMtrCAB-low gave the largest dynamic range in cytochrome expression and was used for all subsequent experiments. To assess the ability for pMtrCAB-low to tune cytochrome expression levels in cells using vanillic acid, we transformed a *S. oneidensis* mutant devoid of outer-membrane cytochromes (ΔMTR) with the pMtrCAB-low plasmid^41^. To determine the relative cytochrome expression levels achievable, we used SDS-PAGE and heme-staining to separate and visualize cytochromes, respectively. *S. oneidensis* ΔMTR + pMtrCAB-low was found to display maximal cytochrome expression using 25 µM vanillic acid (**Figure 2B**). *S. oneidensis* ΔMTR transformed with an empty vector only showed the periplasmic and inner-membrane cytochromes FccA (63 kDa) and CymA (21 kDa), respectively, as expected^41^. Additionally, to quantify the abundance of the MtrCAB conduit on the cell surface we generated pMtrCAB-low-ST where MtrC contains a SpyTag peptide insertion following amino acid 245 and labelled cell surface MtrC using SpyCatcher-GFP^42^. Following growth with varying vanillic acid concentrations, we observed that cytochrome concentration could be tuned between 1.5 ± 0.7 x 10^3^ MtrC cell^-1^ in the absence of vanillic acid to 1.0 ± 0.04 x 10^5^ MtrC cell^-1^ in the presence of 25 μM vanillic acid (**Figure S2, Table S3**). The cytochrome concentration when induced with 25 μM vanillic acid is in close agreement with estimates of cytochrome density from electron cryotomography^43^. Thus, pMtrCAB-low in *S. oneidensis* ΔMTR enabled chemogenetic control over extracellular cytochrome expression over 2 orders-of-magnitude.

We next assessed if a strain containing both the optogenetic patterning and chemical-inducible cytochrome plasmids could be patterned with light and exhibit tunable EET as a function of added vanillic acid. To determine if EET activity could be controlled by tuning cytochrome concentration, we measured the electrochemical activity of wild-type *S. oneidensis* MR-1, *S. oneidensis* ΔMTR, and *S. oneidensis* ΔMTR + pMtrCAB-low, all containing the optogenetic patterning plasmid. Biofilms of fixed surface area (1.2 cm x 1 cm) were patterned onto transparent indium-tin oxide (ITO) electrodes by culturing cells in custom, glass tube reactors under blue-light illumination.

Cyclic voltammetry was then performed in the presence of the electron donor lactate. WT *S. oneidensis* generated a catalytic current at potentials >0.1 V_SHE_ while ΔMTR did not (**Figure 2C**). Thus, we observed the expected defect in extracellular electron transfer with the absence of surface cytochromes. ΔMTR transformed with pMtrCAB- low showed an increase in catalytic current in the presence of increasing concentrations of vanillic acid (**Figure 2D**). The maximum current generated from ΔMTR + pMtrCAB- low reached the same level as the WT *S. oneidensis,* suggesting that this plasmid expresses cytochromes at a comparable level to WT. Collectively, these experiments illustrate that this strain can be optogenetically patterned with independent control of EET activity using vanillic acid.

### Cytochrome doping enables tunable conductivity in patterned biofilms

In addition to enabling heterogenous electron transfer between biofilms and electrodes, extracellular cytochromes also facilitate long-distance electron transport through biofilms via homogenous electron transfer between cytochromes. This long-distance electron transport enables cells that are not directly in contact with electrodes to couple their metabolism at a distance using extracellular cytochromes^44^. To determine how the concentration of outer-membrane cytochromes influences biofilm conductivity, we used electrochemical gating to measure the conductivities of patterned biofilms doped with varying cytochrome content. Electrochemical gating, originally developed for measuring electron transport in conductive polymers^45,46^, uses interdigitated electrodes (IDEs) to measure charge transport through thin films and has previously been used to measure biofilm conduction^9,10,12,47^. In electrochemical gating, a sample is deposited onto two IDEs, with one electrode serving as a source electrode and the other serving as a drain electrode (**Figure 3A-B**). Then, similarly to cyclic voltammetry, the potential at each electrode is swept across a defined range, now with a gate electrode being used to create a small potential offset (*V*_SD_) between the source and drain electrodes. The current flow between the two electrodes, *I*_cond_ (conduction current), is then measured as the gate potential (*E*_gate_) is varied. Here, *E*_gate_ is the average of the potential at the source (*E*_S_) and drain (*E*_D_) electrodes. For redox conduction, as exhibited by conductive biofilms, as *E*_gate_ approaches the redox potential of the charge carrying cytochromes, current flows from one electrode to the other, through the biofilm as the cytochrome exist in a mixed population of oxidized and reduced states^9,47^. However, when *E*_gate_ diverges and the Fermi levels of the electrodes become misaligned with the redox potential of the cytochromes, homogenous cytochrome-cytochrome electron transfer becomes limited by depletion of either the reduced or oxidized population eliminating the ability for a redox gradient to form. This limits the number of electrons that can enter the biofilm and is observed as a peak-shaped dependence of *I*_cond_ on *E*_gate_.

Electrochemical gating measurements performed on patterned ΔMTR + pMtrCAB- low biofilms at varying vanillic acid concentrations showed the expected peak-shaped dependence of *I*_cond_ on *E*_gate_ (**Figure 3C**). Similar to the trend observed in our ΔMTR + pMtrCAB-low cyclic voltammetry measurements, we see each *I*_cond_ peak increasing with increasing vanillic acid concentration. Here, the different *I*_cond_ peaks remain centered at +0.18 V_SHE_, near the formal redox potential of MtrC^9,48^. Interestingly, the *I*_cond_ observed at the maximum vanillic acid concentration used was similar to that of WT *S. oneidensis*, with the WT redox peak shifted ∼25 mV more positive to +0.2 V_SHE_ (**Figure S3**). This again suggests that WT *S. oneidensis* may already possess optimized cytochrome expression levels and that the expression of alternative surface cytochromes (*e.g.,* OmcA, MtrF) impacts the observed redox features. Additionally, the changes in electrochemical activity and in conduction observed in response to changes in vanillic acid further confirm that the MtrCAB complex mediates homogenous (cell-cell) electron transport in biofilm conduction, in addition to heterogenous (electrode-cell) electron transport.

To derive the biofilm conductivity (σ) for each induction conditions, we used Equations 1-3 (Methods) and the the measured peak *I*_cond_, the known *V*_SD_, the known biofilm geometries obtained from light patterning, and the median biofilm thickness as measured across reactors (10 µm, **Figure S4**). Additionally, as the expression used relies on a linear relationship between *I*_cond_ and *V*_SD_, we confirmed that across the *V*_SD_ potential range of 10 – 60 mV, *S. oneidensis* WT displayed the expected peak-shaped *I*_cond_ centered at the redox potential of the cell surface cytochromes and it displayed a peak *I*_cond_ linearly proportional to *V*_SD_^48^ (**Figure S5**, r^2^ = 0.991). With this, we determined the dynamic range of biofilm conductivity for our system to be 1.3 – 6.3 nS cm^-1^ at the varying MtrC concentrations on the cell surface used here (**Figure 3D**). Based on a Hill function fit to the measured conductivities, half-maximal biofilm conductivity is achieved at an MtrC concentration of 3.3 x 10^4^ MtrC cell^-1^, with concentrations above 4.8 x 10^4^ MtrC cell^-1^ yielding similar conductivities to that of the maximally induced condition (∼6 nS cm^-1^). Collectively these results show that by doping living biofilms with different concentrations of cell surface cytochromes, we can tune their intrinsic electronic conductivity. Additionally, with cell surface cytochromes under the control of a small molecule inducible promoter, these biofilms behave as living chemiresistors, changing the biofilm conductivity and the current capable of flowing through the biofilm in response to the presence of different vanillic acid concentrations.

### Thermal dependence of electron transport in biofilms depends on cytochrome concentration

To reveal mechanistic information about how cell surface cytochrome concentration impacts biofilm conduction, we determined the electron transport activation energies (*E*_A_) at each vanillic acid concentration. Here, activation energy represents the effective energy barrier required for biofilm conduction, encompassing the different physical mechanisms that enable this conduction. To calculate activation energies, we performed electrochemical gating measurements on patterned biofilms of ΔMTR + pMtrCAB-low at four different temperatures (28°C, 31°C, 34°C, and 38.5°C) for each vanillic acid concentration and determined the corresponding biofilm conductivities (**Figure 4A**). The biofilms were assumed to behave as ideal redox conductors, and the resulting peak *I*_cond_ values obtained at each temperature point were fit with an Arrhenius relationship to determine the activation energy for each vanillic acid concentration (**Figure S6**).

**Figure 4.**
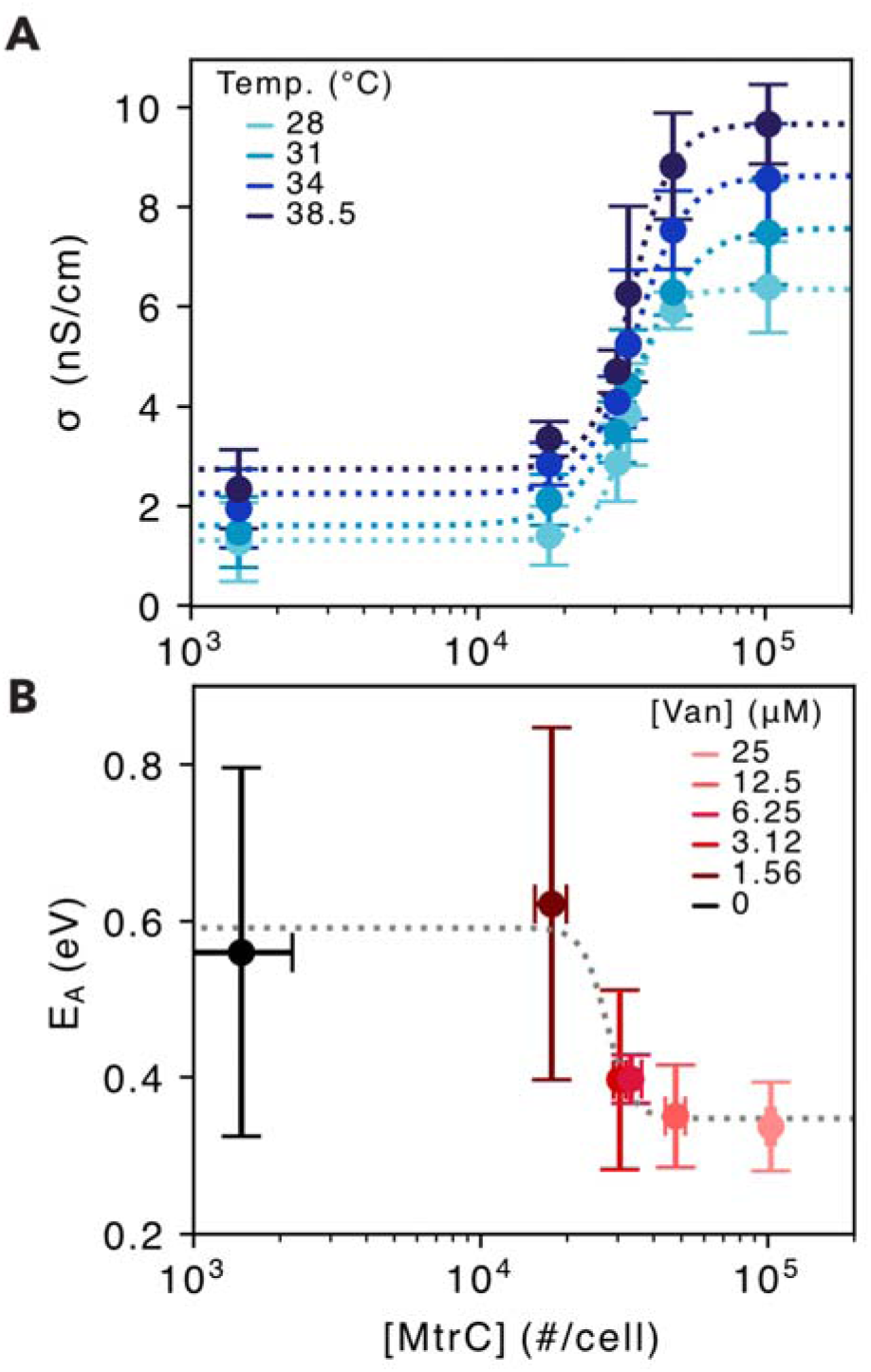
Temperature dependence of biofilm conductivity with varying cytochrome concentrations. **(A)** Biofilm conductivity relative to the cytochrome abundance per cell measured at varying temperatures **(B)** Thermal activation energy of biofilms with varying concentrations of cytochromes. Circles and error bars represent the mean and standard deviation, respectively, of biological replicates (n=3-5 per condition). Dotted lines represent Hill function fits to the data to calculate the half- maximal cytochrome concentrations.

From the data, we see a modest trend of decreasing activation energy as the cytochrome concentration increases. Interestingly, as we approached the maximum cytochrome concentration, 1.0 ± 0.04 x 10^5^ MtrC cell^-1^, we see the activation energy (0.32 ± 0.04 eV) approaches the activation energies for conduction in light-patterned WT *S. oneidensis* MR-1 (0.29 ± 0.08 eV**, Figure S7**) as well as values previously reported for non-patterned WT *S. oneidensis*^9,49^ and for intermolecular electron transfer between MtrC cytochromes^50^ (**Figure 4B**). As cytochrome concentration decreases, there is an increase in the activation energy to 0.54 ± 0.04 eV. Proteins, including the multiheme cytochromes responsible for mediating EET and conduction in electroactive biofilms, can diffuse laterally along the surface of the outer membrane^51^. It is also known that protein crowding on the outer membrane surface can lower the diffusion coefficients of these proteins^52^. Because cytochromes can diffuse along cell surfaces, biofilm electron transport can be modeled as a mix of electron hopping and cytochrome diffusion. This has been dubbed the collision-exchange model of electron transport as it considers cytochrome-cytochrome collisions resulting from their movement along the cell surface as well as resulting from intermolecular electron exchanges occurring between colliding cytochromes^43,51^. One explanation for the increased activation energies is that as cytochrome expression was lowered, conduction enters a regime where lateral cytochrome diffusion limits electron transfer due to an increased time between cytochrome-cytochrome interactions as previously suggested by Monte Carlo simulations^51^. Here, we demonstrate that in addition to bioelectronics applications, tuning cytochrome concentrations offers new mechanistic insights into the regime and limits of biological electron transport.

### Living electronic films display negative differential transconductance

In addition to their chemiresistive properties, these engineered biofilms can function as living transistors with the conductive biofilms acting as gateable channels spanning source-drain electrodes. In this case, it is the electronic structure of the cytochromes that enables charge transfer, but only when the potentials applied to the source-drain electrodes are near the redox potential of the cytochromes. Unlike traditional solid-state field-effect transistors (FETs), or organic electrochemical transistors (OECTs), the conduction current does not solely increase (n-type FET or accumulation-mode OECT) or decrease (p-type or depletion-mode OECT) with increasing gate potential^53,54^. The peak shaped relationship between current (*I*_cond_) and gate potential (*E*_gate_) observed for conductive biofilms is similar to that of an anti-ambipolar transistor (AAT), despite stemming from a different mechanism. AATs are built with an anti-ambipolar heterojunction channel consisting of overlapping n-type and p-type material that each span partway across the source-drain electrode gap resulting in peak shaped conductance due to the overlaps in electron and hole transport^55,56^. To better relate the *I*_cond_ vs *E*_gate_ characteristics of living electronic films to those of solid-state AATs, we have calculated the differential transconductance (g_m_) from our electrochemical gating data and we extracted common AAT figures of merit (**Figure 5A-B**). The living electronic films displayed the characteristic negative differential transconductance (NDT) that is a hallmark of anti-ambipolarity (**Figure 5C**). At the maximum cytochrome concentration (1.0 ± 0.04 x 10^5^ MtrC cell^-1^) and with *V*_SD_ = 20 mV, the living electronic films displayed a peak conductance (*I*_peak_) of ∼22 ± 1.4 nA, a peak voltage (*V*_peak_) at *E*_gate_ = 0.184 ± 0.002 V_SHE_, a peak-to-valley (PVR = *I*_peak_/*I*_valley_) ratio of 49 ± 11.1, and an on voltage (V_ON_), or peak width, of 0.227 ± 0.007 V. The WT *S. oneidensis* biofilms yielded similar AAT behaviors (**Figure S8**). Relative to heterojunction AAT devices, living electronic films operate in a low power regime dictated by thermodynamic redox potentials of the cytochromes. This results in a sharp slope in the transconductance over hundreds of millivolts, whereas heterojunction AAT devices typical require steps spanning tens of volts to generate NDT^56^. AATs and other NDT electronics show promise with applications in frequency modulation, neuromorphic computing, and multivalued logic devices^56^, suggesting that living electronic films exhibiting NDT behavior could find future applications in similar areas.

**Figure 5.**
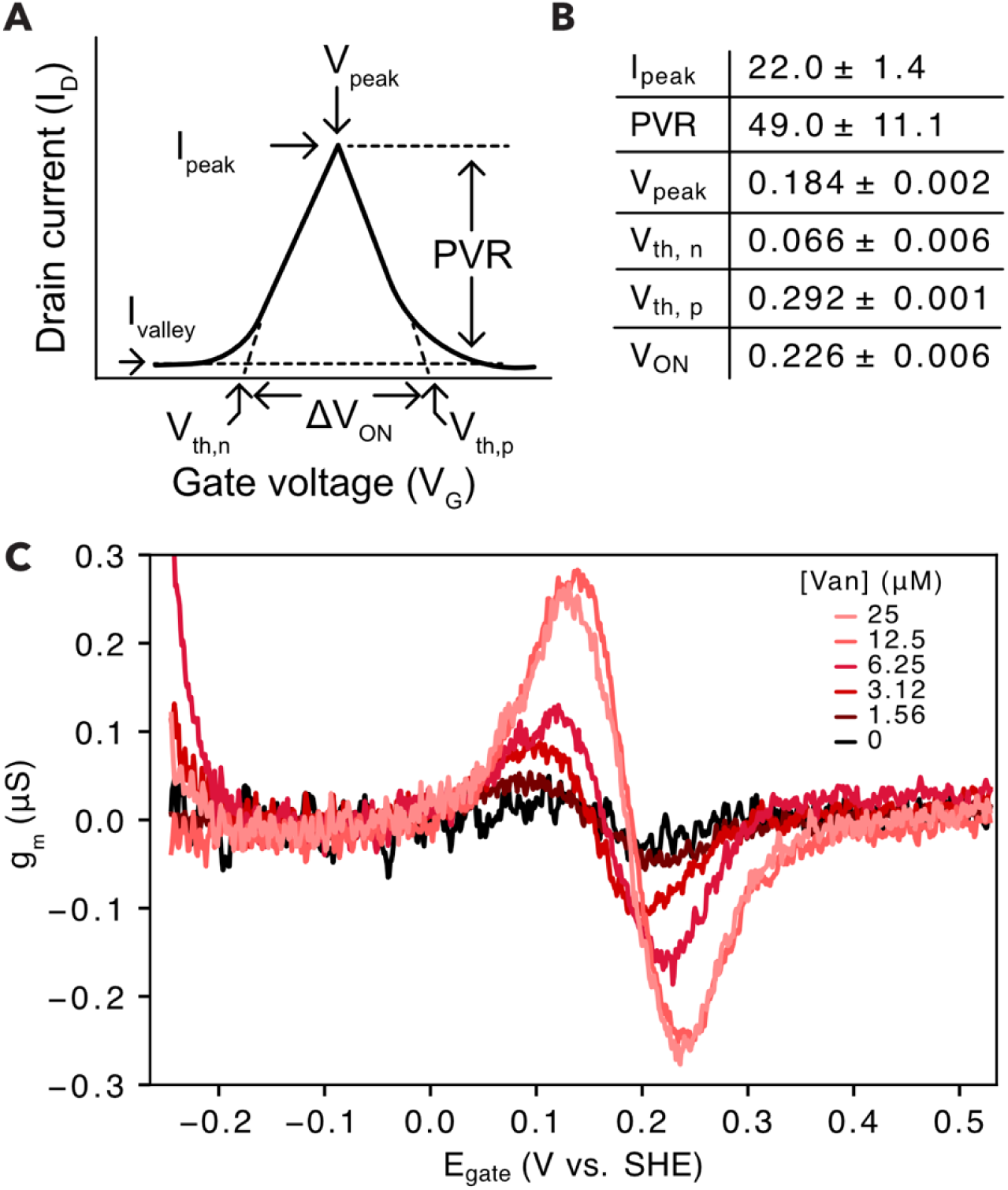
Biofilm differential transconductance and anti-ambipolar transistor figures of merit. **(A)** Characteristic I_cond_ vs E_gate_ behavior for anti-ambipolar transistors (AATs) and other negative differential transconductance (NDT) devices. Feature labels highlight figures of merit typically extracted from this data^56^. **(B)** AAT figures of merit extracted from the I_cond_ vs E_gate_ data for the ΔMTR containing the pMtrCAB-low genetic circuit and in the presence of 25 μM vanillic acid from Figure 3C. **(C)** Differential transconductance (g_m_), obtained from taking the derivative of conduction current (I_cond_) with respect to gate potential (E_gate_), plotted against gate potential (E_gate_) for patterned biofilms of the ΔMTR + pMtrCAB-low in the presence of various concentrations of vanillic acid. Data from the same replicates used in Figure 3C are used here.

## CONCLUSION

In summary, we utilized synthetic biology techniques to simultaneously control the deposition and electrical properties of conductive, *S. oneidensis* biofilms. This was achieved by modifying cells with both the optogenetic patterning gene circuit and a chemogenetic gene circuit to control cytochrome expression with a small molecule. Electrochemical activity and gateable biofilm conduction showed that conductivity could be tuned over a 6-fold range by varying cell surface cytochromes by 2 orders-of-magnitude. Biofilm conductivity for patterned WT *S. oneidensis* was found to be similar in magnitude to that of the maximally induced strain, 5.74 ± 0.20 nS cm^-1^ and 6.38 ± 0.91 nS cm^-1^, respectively, suggesting that WT *S. oneidensis* has natively maximized cytochrome expression. Additionally our cytochrome labelling experiments showed ΔMTR + pMtrCAB-low when optimally induced produced 1.0 ± 0.04 x 10^5^ MtrC cell^-1^, which is in close agreement with estimates of cytochrome density from electron cryotomography^43^. The temperature dependence of conduction revealed that the activation energy is dependent on cytochrome concentration, suggesting a transition between multiple mechanistic regimes. At high cytochrome concentrations the activation energy approached that of predicted activation energies for intermolecular electron transfer between cytochromes, suggesting that the cells have entered a percolation regime where electron transport is primarily limited by electron transfer between cytochromes. By contrast, an increase in activation energy was observed for decreasing cytochrome concentrations suggesting electron transport is limited by another physical process, likely cytochrome diffusion.

With biofilm conduction under the direct control of vanillic acid, we have effectively created a living, self-assembled chemiresistor. By coupling cytochrome expression to chemogenetic promoters that respond to other analytes, this system has the potential to be used as a modular biosensing platform for diverse analytes. This work has defined essential design principles governing chemiresistance in *S. oneidensis* biofilms, specifically the cytochrome concentrations regimes needed to tune biofilm conductivity. In addition to their chemiresistive properties, these engineered biofilms function as electrolyte-gated transistors with a biological semiconducting layer where current can be controlled by a gate electrode and is set by the reduction potential and concentration of the outer membrane cytochromes in the biofilm^57^. These systems could be expanded to respond to varying gate potentials by modifying which cytochrome is expressed on the cell surface or coupling this with de novo design to modulate cytochrome redox potentials^58^. Thus, with this system, we lay the foundations for the construction of living electronic devices and have created a platform for examining fundamental mechanisms of conduction in electroactive biofilms.

## ACKNOWLEDGEMENTS

pAJM.773 was a gift from Christopher Voigt (Addgene plasmid #108527). pSpyCatcher003-sfGFP was a gift from Mark Howarth (Addgene plasmid #133449). The fabrication of the ITO-IDEs used in this work was performed at the San Diego Nanotechnology Infrastructure (SDNI) of UCSD, a member of the National Nanotechnology Coordinated Infrastructure, which is supported by the National Science Foundation.

## DATA AVAILABILITY

Designs for custom IDE devices can be found in a Zenodo repository (10.5281/zenodo.21959616). Electrochemistry data and scripts for analysis and figure generation are available as supplemental data. All other data, plasmids, and strains are available from the corresponding author on reasonable request.

## REFERENCES

1. Myers, C. R. & Nealson, K. H. Bacterial manganese reduction and growth with manganese oxide as the sole electron acceptor. Science 240, 1319–1321 (1988).

2. Lovley, D. R., Stolz, J. F., Nord, G. L. & Phillips, E. J. P. Anaerobic production of magnetite by a dissimilatory iron-reducing microorganism. Nature 330, 252–254 (1987).

3. Shi, L. et al. Extracellular electron transfer mechanisms between microorganisms and minerals. Nat. Rev. Microbiol. 14, 651–662 (2016).

4. Edwards, M. J. et al. Structural modeling of an outer membrane electron conduit from a metal-reducing bacterium suggests electron transfer via periplasmic redox partners. J. Biol. Chem. 293, 8103–8112 (2018).

5. Fonseca, B. M. et al. Mind the gap: cytochrome interactions reveal electron pathways across the periplasm of Shewanella oneidensis MR-1. Biochem. J 449, 101–108 (2013).

6. Hartshorne, R. S. et al. Characterization of an electron conduit between bacteria and the extracellular environment. Proc. Natl. Acad. Sci. U. S. A. 106, 22169– 22174 (2009).

7. White, G. F. et al. Rapid electron exchange between surface-exposed bacterial cytochromes and Fe(III) minerals. Proc. Natl. Acad. Sci. U. S. A. 110, 6346–6351 (2013).

8. Edwards, M. J., White, G. F., Butt, J. N., Richardson, D. J. & Clarke, T. A. The crystal structure of a biological insulated transmembrane molecular wire. Cell 181, 665–673.e10 (2020).

9. Xu, S., Barrozo, A., Tender, L. M., Krylov, A. I. & El-Naggar, M. Y. Multiheme Cytochrome Mediated Redox Conduction through Shewanella oneidensis MR-1 Cells. J. Am. Chem. Soc. 140, 10085–10089 (2018).

10. Yates, M. D. et al. Thermally activated long range electron transport in living biofilms. Phys. Chem. Chem. Phys. 17, 32564–32570 (2015).

11. Snider, R. M., Strycharz-Glaven, S. M., Tsoi, S. D., Erickson, J. S. & Tender, L. M. Long-range electron transport in Geobacter sulfurreducens biofilms is redox gradient-driven. Proc. Natl. Acad. Sci. U. S. A. 109, 15467–15472 (2012).

12. Zhao, F. et al. Light-induced patterning of electroactive bacterial biofilms. ACS Synth. Biol. 11, 2327–2338 (2022).

13. Logan, B. E. & Rabaey, K. Conversion of wastes into bioelectricity and chemicals by using microbial electrochemical technologies. Science 337, 686–690 (2012).

14. Zacharoff, L. A. & El-Naggar, M. Y. Redox conduction in biofilms: From respiration to living electronics. Curr. Opin. Electrochem. 4, 182–189 (2017).

15. Atkinson, J. T., Chavez, M. S., Niman, C. M. & El-Naggar, M. Y. Living electronics: A catalogue of engineered living electronic components. Microb. Biotechnol. 16, 507–533 (2023).

16. Zhang, Y., Hsu, L. H.-H. & Jiang, X. Living electronics. Nano Res. 13, 1205–1213 (2020).

17. Sun, J. et al. Living synthelectronics: A New Era for bioelectronics powered by synthetic biology. Adv. Mater. 2400110 (2024).

18. Shi, J. et al. Active biointegrated living electronics for managing inflammation. Science 384, 1023–1030 (2024).

19. Levskaya, A. et al. Synthetic biology: engineering Escherichia coli to see light: Synthetic biology. Nature 438, 441–442 (2005).

20. Piraner, D. I., Abedi, M. H., Moser, B. A., Lee-Gosselin, A. & Shapiro, M. G. Tunable thermal bioswitches for in vivo control of microbial therapeutics. Nat. Chem. Biol. 13, 75–80 (2017).

21. Meyer, A. J., Segall-Shapiro, T. H., Glassey, E., Zhang, J. & Voigt, C. A. Escherichia coli "Marionette’’ strains with 12 highly optimized small-molecule sensors. Nat. Chem. Biol. 15, 196–204 (2019).

22. Tschirhart, T. et al. Electronic control of gene expression and cell behaviour in Escherichia coli through redox signalling. Nat. Commun. 8, 14030 (2017).

23. Bird, L. J. et al. Engineering wired life: Synthetic biology for electroactive bacteria. ACS Synth. Biol. 10, 2808–2823 (2021).

24. Jensen, H. M. et al. Engineering of a synthetic electron conduit in living cells. Proc. Natl. Acad. Sci. U. S. A. 107, 19213–19218 (2010).

25. Sturm-Richter, K. et al. Unbalanced fermentation of glycerol in Escherichia coli via heterologous production of an electron transport chain and electrode interaction in microbial electrochemical cells. Bioresour. Technol. 186, 89–96 (2015).

26. Jensen, H. M., TerAvest, M. A., Kokish, M. G. & Ajo-Franklin, C. M. CymA and exogenous flavins improve extracellular electron transfer and couple it to cell growth in Mtr-expressing Escherichia coli. ACS Synth. Biol. 5, 679–688 (2016).

27. West, E. A., Jain, A. & Gralnick, J. A. Engineering a Native Inducible Expression System in Shewanella oneidensis to Control Extracellular Electron Transfer. ACS Synth. Biol. 6, 1627–1634 (2017).

28. Simoska, O., Gaffney, E. M., Lim, K., Beaver, K. & Minteer, S. D. Understanding the Properties of Phenazine Mediators that Promote Extracellular Electron Transfer in Escherichia coli. J. Electrochem. Soc. 168, 025503 (2021).

29. Wu, Z. et al. Engineering an electroactive Escherichia coli for the microbial electrosynthesis of succinate from glucose and CO2. Microb. Cell Fact. 18, 15 (2019).

30. Su, L. et al. Modifying Cytochrome c Maturation Can Increase the Bioelectronic Performance of Engineered Escherichia coli. ACS Synth. Biol. 9, 115–124 (2020).

31. Su, L., Fukushima, T. & Ajo-Franklin, C. M. A hybrid cyt c maturation system enhances the bioelectrical performance of engineered Escherichia coli by improving the rate-limiting step. Biosens. Bioelectron. 165, 112312 (2020).

32. Long, X., Tokunou, Y. & Okamoto, A. Mechano-control of extracellular electron transport rate via modification of inter-heme coupling in bacterial surface cytochrome. Environ. Sci. Technol. (2023) doi:10.1021/acs.est.3c00601.

33. Mouhib, M., Reggente, M., Li, L., Schuergers, N. & Boghossian, A. A. Extracellular electron transfer pathways to enhance the electroactivity of modified Escherichia coli. Joule 7, 2092–2106 (2023).

34. Zhao, F. et al. Red-light-induced genetic system for control of extracellular electron transfer. ACS Synth. Biol. 13, 1467–1476 (2024).

35. Zhao, F. et al. Light-directed biofilm formation reveals the functional contributions of periplasmic cytochromes to the electrochemical activity of *Shewanella oneidensis*. MBio (2026) doi:10.1128/mbio.01264-26.

36. Engler, C., Kandzia, R. & Marillonnet, S. A one pot, one step, precision cloning method with high throughput capability. PLoS One 3, e3647 (2008).

37. Reis, A. C. & Salis, H. M. An automated model test system for systematic development and improvement of gene expression models. ACS Synth. Biol. 9, 3145–3156 (2020).

38. Bretschger, O. et al. Current production and metal oxide reduction by Shewanella oneidensis MR-1 wild type and mutants. Appl. Environ. Microbiol. 73, 7003–7012 (2007).

39. Kieft, T. L. et al. Dissimilatory reduction of Fe(III) and other electron acceptors by a Thermus isolate. Appl. Environ. Microbiol. 65, 1214–1221 (1999).

40. Gross, B. J. & El-Naggar, M. Y. A combined electrochemical and optical trapping platform for measuring single cell respiration rates at electrode interfaces. Rev. Sci. Instrum. 86, 064301 (2015).

41. Coursolle, D. & Gralnick, J. A. Reconstruction of extracellular respiratory pathways for iron(III) reduction in Shewanella oneidensis strain MR-1. Front. Microbiol. 3, 56 (2012).

42. Keeble, A. H. et al. Approaching infinite affinity through engineering of peptide-protein interaction. Proc. Natl. Acad. Sci. U. S. A. 116, 26523–26533 (2019).

43. Subramanian, P., Pirbadian, S., El-Naggar, M. Y. & Jensen, G. J. Ultrastructure of Shewanella oneidensis MR-1 nanowires revealed by electron cryotomography. Proc. Natl. Acad. Sci. U. S. A. 115, E3246–E3255 (2018).

44. Pirbadian, S., Chavez, M. S. & El-Naggar, M. Y. Spatiotemporal mapping of bacterial membrane potential responses to extracellular electron transfer. Proc. Natl. Acad. Sci. U. S. A. 117, 20171–20179 (2020).

45. Paul, E. W., Ricco, A. J. & Wrighton, M. S. Resistance of polyaniline films as a function of electrochemical potential and the fabrication of polyaniline-based microelectronic devices. J. Phys. Chem. 89, 1441–1447 (1985).

46. Chidsey, C. E. & Murray, R. W. Electroactive polymers and macromolecular electronics. Science 231, 25–31 (1986).

47. Yates, M. et al. Characterizing electron transport through living biofilms. J. Vis. Exp. (2018) doi:10.3791/54671.

48. Xu, S., Jangir, Y. & El-Naggar, M. Y. Disentangling the roles of free and cytochrome-bound flavins in extracellular electron transport from Shewanella oneidensis MR-1. Electrochim. Acta 198, 49–55 (2016).

49. Wen, X., Long, X., Huang, W., Kuramochi, M. & Okamoto, A. Cell-surface inter- cytochrome electron transfer limits biofilm electron conduction kinetics in Shewanella oneidensis. J. Am. Chem. Soc. (2025) doi:10.1021/jacs.5c10357.

50. Breuer, M., Rosso, K. M., Blumberger, J. & Butt, J. N. Multi-haem cytochromes in Shewanella oneidensis MR-1: structures, functions and opportunities. J. R. Soc. Interface 12, 20141117 (2015).

51. Chong, G. W. et al. Single molecule tracking of bacterial cell surface cytochromes reveals dynamics that impact long-distance electron transport. Proc. Natl. Acad. Sci. U. S. A. 119, e2119964119 (2022).

52. Javanainen, M., Martinez-Seara, H., Metzler, R. & Vattulainen, I. Diffusion of integral membrane proteins in protein-rich membranes. J. Phys. Chem. Lett. 8, 4308–4313 (2017).

53. Rivnay, J. et al. Organic electrochemical transistors. Nat. Rev. Mater. 3, 1–14 (2018).

54. Neamen, D. A. Semiconductor Physics And Devices. (McGraw-Hill Education, 2011).

55. Jariwala, D. et al. Gate-tunable carbon nanotube-MoS2 heterojunction p-n diode. Proc. Natl. Acad. Sci. U. S. A. 110, 18076–18080 (2013).

56. Meng, Y. et al. Anti-ambipolar heterojunctions: Materials, devices, and circuits. Adv. Mater. 36, e2306290 (2024).

57. Torricelli, F. et al. Electrolyte-gated transistors for enhanced performance bioelectronics. Nat. Rev. Methods Primers 1, 1–24 (2021).

58. Hardy, B. J. et al. Cellular production of a de novo membrane cytochrome. Proc. Natl. Acad. Sci. U. S. A. 120, e2300137120 (2023).

