## Supplemental Information for "Living electronic transistors with tunable conductivity"

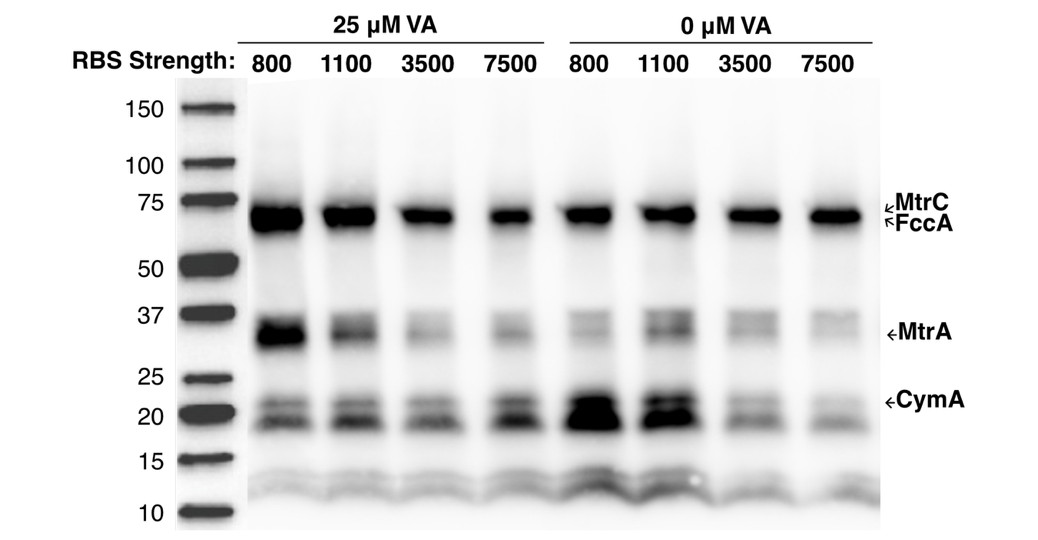

**Figure S1.** Comparison of cytochrome expression from plasmids with varying RBS strengths. An electrochemiluminsence blot of cultures of a multiheme cytochrome mutant (ΔMTR) with plasmids pMtrCAB-low, pMtrCAB-med, pMtrCAB-high, or pMtrCAB-highest that contain RBS with varying translation initiation rates of 800, 1100, 3500, or 7500, respectively, in the presence or absence of 25 μM vanillic acid (VA). MtrA and MtrC bands increase for pMtrCAB-low and pMtrCAB-med in the presence of VA relative to the absence of VA. pMtrCAB-high and pMtrCAB-highest do not appear to express functional MtrC or MtrA in the presence or absence of VA. FccA, a genomically-encoded cytochrome is present under all conditions as a loading control.

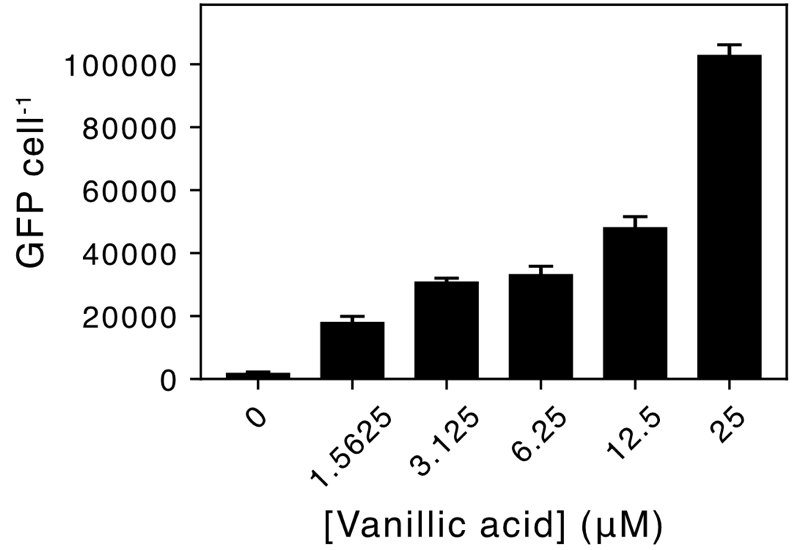

**Figure S2.** SpyCatcher-GFP labelling of MtrC-ST. **(A)** Cells expressing MtrC-ST were labelled with 20 uM of SpyCatcher-GFP and the GFP cell^-1^ was calculated by generating a GFP standard curve and subtracting GFP that non-specifically bound to cell expressing WT MtrC. The bound GFP cell^-1^ was used as a proxy for the surface concentration of MtrC.

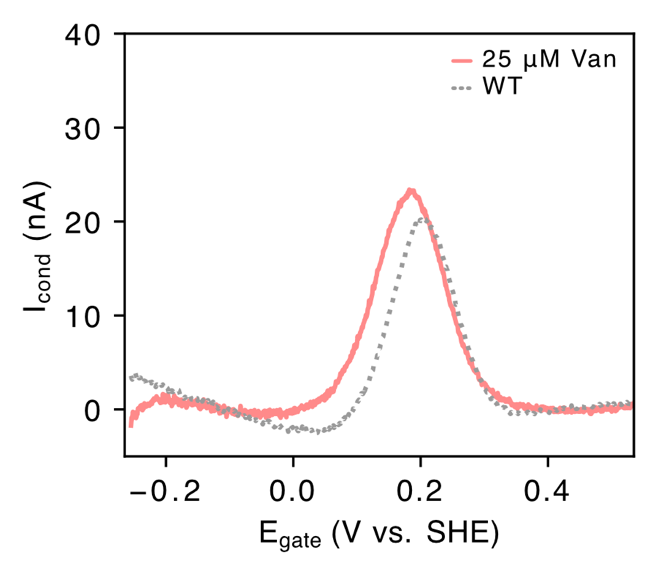

**Figure S3.** Electrochemical gating of electron transport through *S. oneidensis* biofilms. Conduction current (I_cond_) measured from electrochemical gating measurements of patterned biofilms of *S. oneidensis* WT + pDAWN-CdrAB.

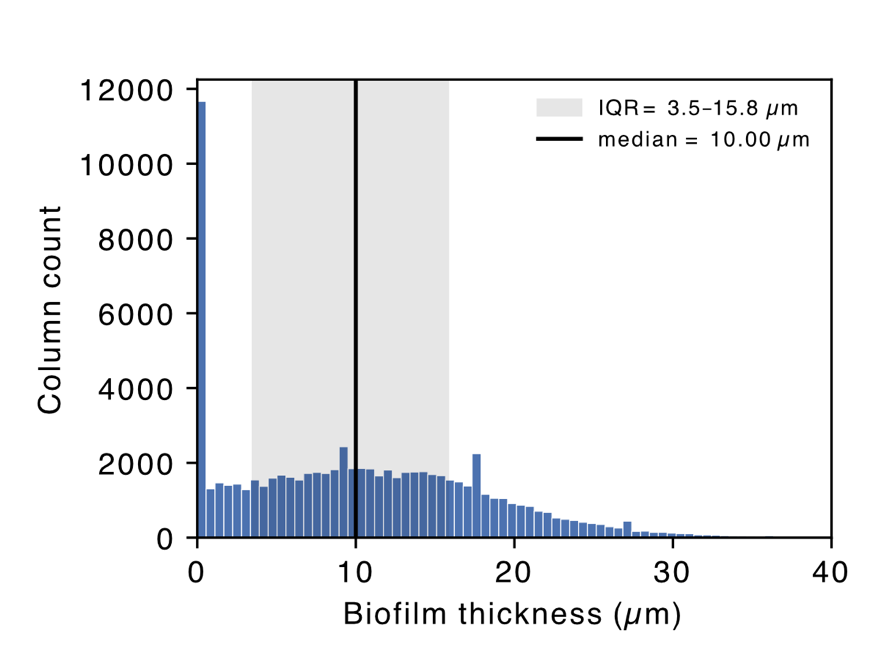

**Figure S4.** Histogram of biofilm thickness scanned across vertical pixel-columns from 50 confocal images from 18 samples. The median biofilm thickness across all columns was 10 μm with an interquartile range of 3.5 – 15.8 μm.

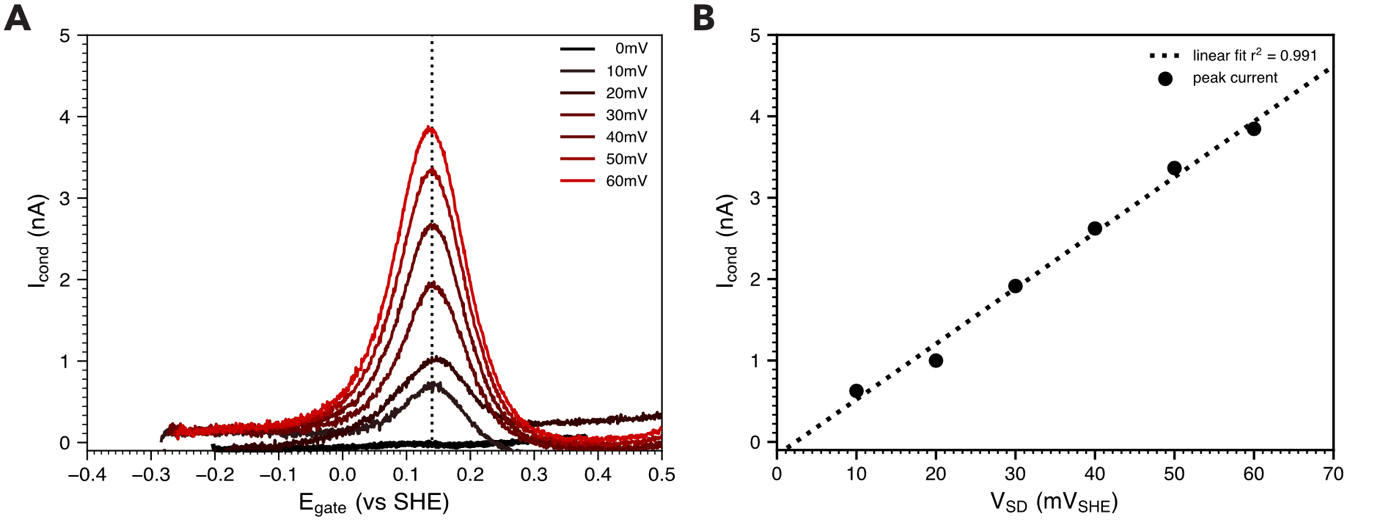

**Figure S5.** Conduction current at varying source-drain voltages (0 – 60 mV). **(A)** Conduction current (I_cond_) measured from electrochemical gating measurements on patterned biofilms of *S. oneidensis* WT + pDAWN-CdrAB with varying varying E_gate_ and V_SD_ on IDEs with 30 um gaps. **(B)** Peak conduction current has a linear response to increased source-drain voltages. Data represent raw conduction currents of a single representative experiment (n=1).

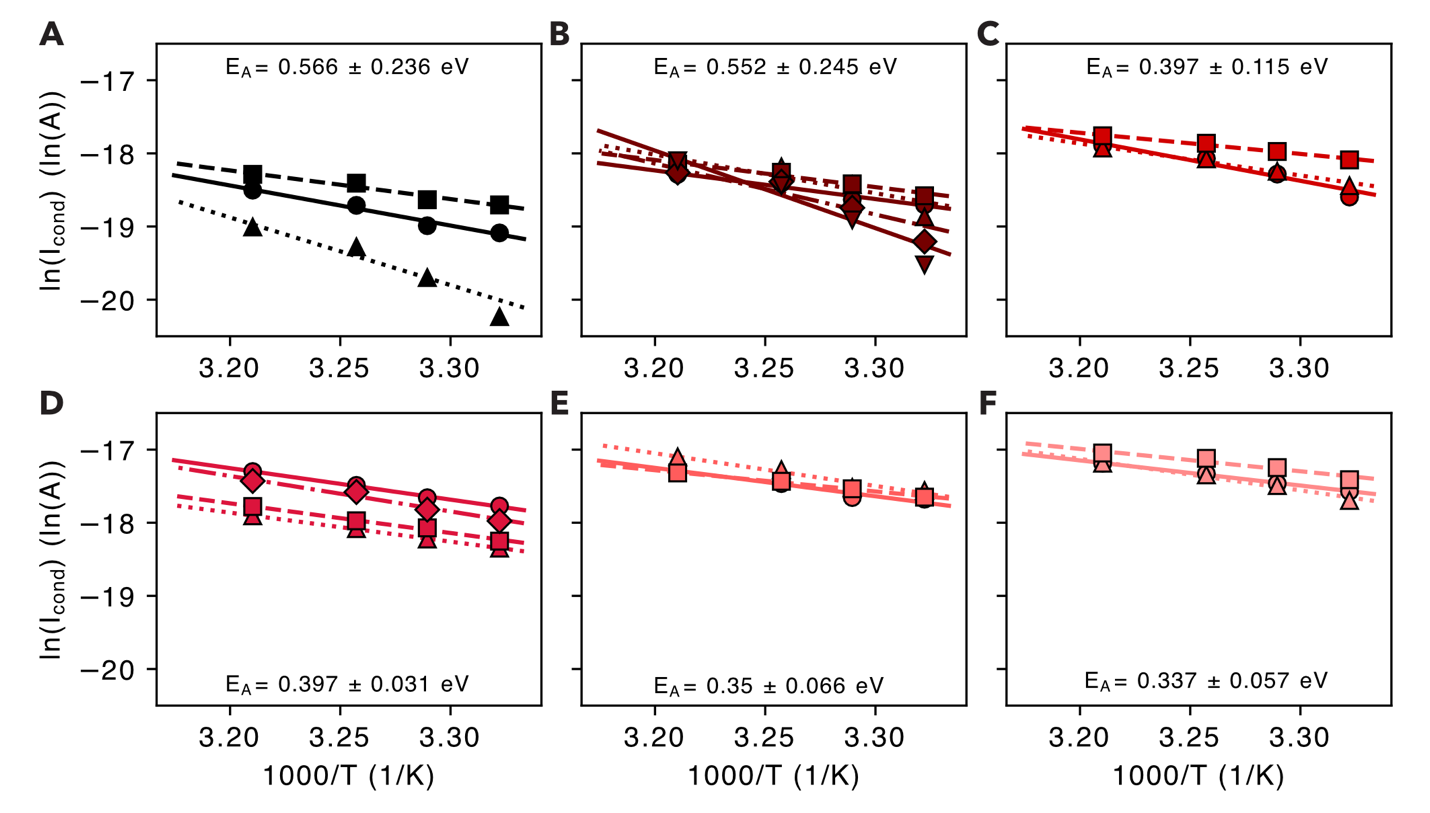

**Figure S6.** Arrhenius plots of ΔMTR + pMtrCAB-low biofilms with varying concentrations of vanillic acid **(A)** 0, **(B)** 1.5625, **(C)** 3.125, **(D)** 6.25, **(E)** 12.5 and **(F)** 25 μM. Individual replicates are plotted as circles, trianges, squares, diamonds, and inverse triangles with linear fit lines. The mean and standard deviation of the activation energy for each concentration is noted (n=3-5 replicates).

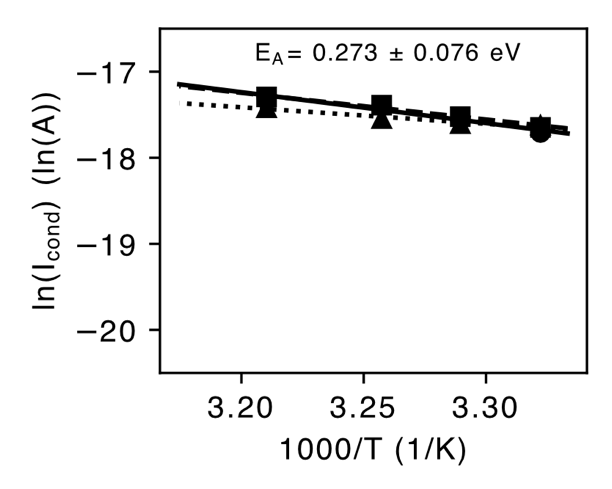

**Figure S7.** Arrhenius plots of *S. oneidensis* WT + pDAWN-CdrAB biofilms. Individual replicates are plotted as circles, triangles, squares. The mean and standard deviation of the activation energy is noted (n=3 replicates).

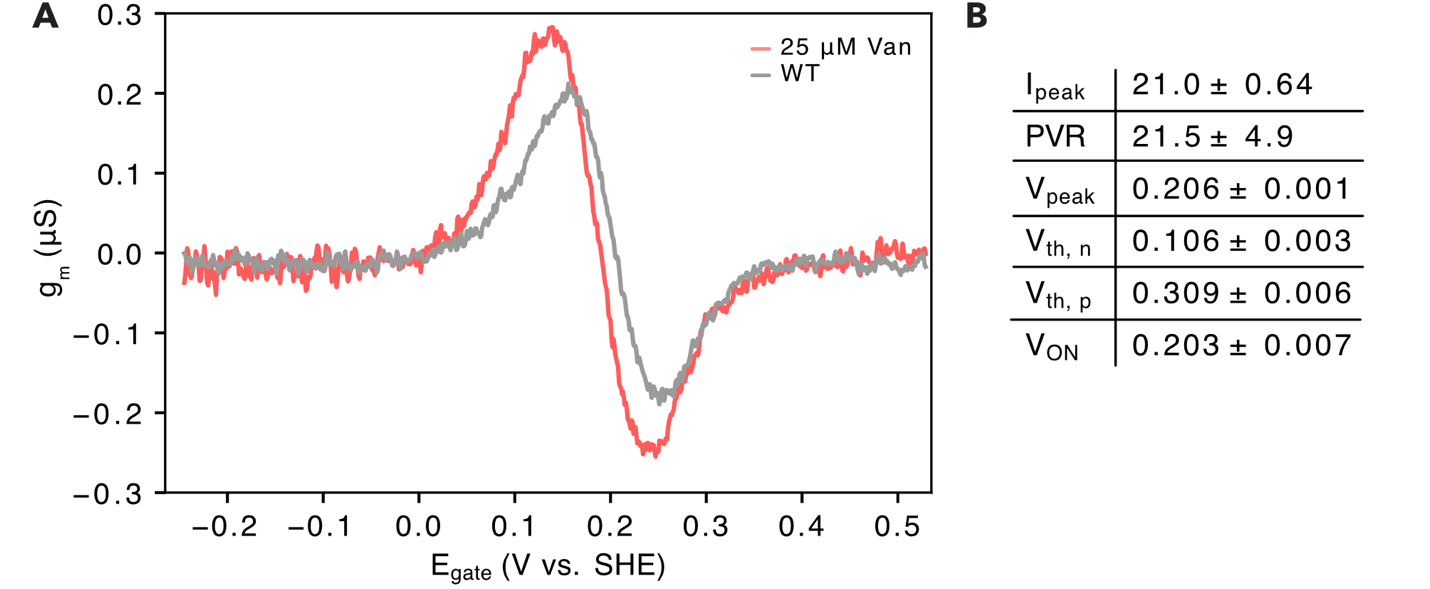

**Figure S8.** Anti-ambipolarity of WT *S. oneidensis* biofilm. **(A)** Transconductance of WT *S. oneidensis* biofilm (gray) relative to ΔMTR + pMtrCAB-low when induced with 25 μM vanillic acid. **(B)** AAT transistor figures-of-merit for *S. oneidensis* WT + pDAWN-CdrAB biofilm.

| **Plasmid** | **Description** | **Source** | **Figure** |
| --- | --- | --- | --- |
| pDAWN-CdrAB | Blue light inducible patterning plasmid, ColE1, *aadA* (Sp^R^) | 12 | 2B, 2D, S1 |
| pAJM.773 | Vector backbone used for generating vanillic acid inducible MtrCAB plasmids,  p15A, *aph(3’)* (Kan^R^) | 21  Addgene (#108527) | - |
| pMtrCAB-low | VanR inducible *mtrCAB* (800 AU RBS),  p15A, *aph(3’)* (Kan^R^) | This work | 2B, 2D, S1 |
| pMtrCAB-med | VanR inducible *mtrCAB* (1100 AU RBS), p15A, *aph(3’)* (Kan^R^) | This work | S1 |
| pMtrCAB-high | VanR inducible *mtrCAB* (3500 AU RBS), p15A, *aph(3’)* (Kan^R^) | This work | S1 |
| pMtrCAB-highest | VanR inducible *mtrCAB* (7500 AU RBS), p15A, *aph(3’)* (Kan^R^) | This work | S1 |
| pMtrCAB-low-ST | VanR inducible *mtrCAB* (800 AU RBS) with SpyTag inserted at amino acid 245 in *mtrC*,  p15A, *aph(3’)* (Kan^R^) | This work | S1 |
| pSpyCatcher003-sfGFP | LacI inducible 6xHis-TEV-SpyCatcher003-sfGFP, colE1, *bla* (AmpR) | 41 Addgene (#133449) | S1 |

**Table S1.** All plasmids used in this study

| **Organism** | **Strain** | **Genotype** | **Source** | **Figure** |
| --- | --- | --- | --- | --- |
| *S. oneidensis* | MR-1 | WT | Lab collection | - |
| *S. oneidensis* | ΔMTR (JG1486) | *ΔmtrB/ΔmtrE/ ΔmtrC/ΔomcA/ ΔmtrF/ΔmtrA/ ΔmtrD/ ΔdmsE/ΔSO4360/ ΔcctA/ ΔrecA* | 40 | - |
| *S. oneidensis* | sJA104 | MR-1 + pDAWN-CdrAB | 12 | 2C, S2 |
| *S. oneidensis* | sJA077 | ΔMTR + pDAWN-CdrAB | This work | 2C, S2 |
| *S. oneidensis* | sJA091 | ΔMTR + pDAWN-CdrAB + pEmpty | This work | 2B |
| *S. oneidensis* | sJA101 | ΔMTR + pDAWN-CdrAB + pMtrCAB-low | This work | S1, 2B, 2D, S2 |
| *S. oneidensis* | sJA102 | ΔMTR + pDAWN-CdrAB + pMtrCAB-med | This work | - |
| *S. oneidensis* | sJA103 | ΔMTR + pDAWN-CdrAB + pMtrCAB-high | This work | **-** |
|  | sJA090 | ΔMTR + pDAWN-CdrAB + pMtrCAB-highest | This work | **-** |
| *E. coli* | BL21(DE3) | *fhuA2 [lon] ompT gal (λ DE3) [dcm] ∆hsdS λ DE3* | NEB | S1 |

**Table S2.** All strains used in this study

| **[Van]**  **(μM)** | **[MtrC] ± SD  (#/cell)** | **Loading Fraction** | **ET rate ± SD**  **(e- s^-1^ cell^-1^)** | **σ ± SD  (nS cm^-1^)** | **E_a_ ± SD  (eV)** |
| --- | --- | --- | --- | --- | --- |
| 0 | 1,471 ± 740 | 0 | 460 ± 234 | 1.28 ± 0.79 | 0.54 ± 0.2 |
| 1.5625 | 17,674 ± 2,219 | 0.014 | 586 ± 184 | 1.41 ± 0.59 | 0.64 ± 0.24 |
| 3.125 | 30,522 ± 1,487 | 0.172 | 1,034 ± 228 | 2.87 ± 0.77 | 0.39 ± 0.1 |
| 6.25 | 32,868 ± 2,951 | 0.326 | 1,385 ± 319 | 3.84 ± 1.02 | 0.4 ± 0.05 |
| 12.5 | 47,775 ± 3,806 | 0.466 | 2,132 ± 109 | 5.92 ± 0.37 | 0.35 ± 0.07 |
| 25 | 102,525 ± 3,665 | 1 | 2,301 ± 269 | 6.38 ± 0.91 | 0.33 ± 0.04 |
| WT | - | - | - | 5.74 ± 0.20 | 0.27 ± 0.08 |

**Table S3.** Summary table of measures and calculated biofilm conductivity parameters.
